# Putting a lid on it: The N-terminal helix of Arf1 inhibits switching via uniform stabilization of the native state

**DOI:** 10.1101/2025.04.22.650101

**Authors:** Edgar V. Peters, Tejaswi Koduru, Noam Hantman, Scott A. McCallum, Qingqiu Huang, Jacqueline Cherfils, Catherine A. Royer

## Abstract

The Arf (and Arf-like) GTPases, unlike all other Ras family GTPase members, exhibit a repressed conformation in the inactive, GDP-bound form. The N-terminal helix of Arf GTPases, which is missing in the other Ras family members, caps the switch region, confining it to this repressed state. Nucleotide exchange and activation involve a massive conformational change made possible by the dissociation of the N-terminal helix from the core of the protein. Spontaneous switching in Arfs is enhanced upon deletion of this helix. While the structural and functional role of the N-terminal helix in Arf proteins is well-known, the energetic basis for its effects have not been established. Here we mapped the local stability of the Arf1Δ17 variant, deleted for the N-terminal helix, using high pressure biophysical approaches and compared it to that of full-length Arf1. Deletion of the N-terminal helix decreased Arf1 stability across the entire structure. Thus, rather than imposing a specific structural pathway for repression, the N-terminal helix exercises global control of Arf1 stability to repress switching.

**Statement of Significance:** Energetic mapping of the N-terminal deletion mutant of Arf1 using high pressure biophysical approaches reveals that this helix maintains the protein in the repressed state unique to Arf and Arf-like GTPases via overall stabilization of the native state rather than unique allosteric communication with the switch region.

## Introduction

ADP ribosylation factor (Arf) family proteins are a subfamily of the Ras family of small GTPases (1). They are implicated in membrane trafficking, motility, division, apoptosis and transcriptional regulation. Arfs are conserved from yeast (2, 3) to humans (4). Because of their central roles in essential cellular processes, Arfs have been implicated in multiple disease states. For example, Arf function is perturbed during viral infection (5, 6). Membrane remodeling and cytoskeleton interactions by Arfs have also been implicated in cancer progression (7, 8). Thus, understanding Arf functional mechanisms and specificities is central to the development of new therapeutic strategies targeting this key family of enzymes.

Small GTPases act as signaling proteins. In their GTP-bound states they are active, transmitting signaling cues via interactions with downstream partners, while when bound by GDP they are inactive. Switching between the inactive to the active state is controlled by dissociation of the GDP, with subsequent binding of the much higher concentration GTP nucleotide, while inactivation requires GTP hydrolysis. Arfs bind GDP very tightly, and require specific guanine nucleotide exchange factors (GEFs) for efficient exchange (9–11). Likewise, Arfs are poor GTP hydrolases, and require GTPase activating proteins (GAPs) to enhance switching to the inactive state (12, 13).

The Arf (and Arf-like) GTPases differ from other Ras family GTPases in that their inactive GDP-bound form exists in a repressed state in which the conformation of the switch region is massively distinct from its structure in the GTP-bound active state, as seen for Arf1 (Figure 1) (14, 15). The N-terminal helix of the Arf proteins acts to constrain the switch region, shifting the interswitch β-hairpin down in register by two residues. In this repressed state, switch 1 forms a β-strand that extends the central β-sheet containing the interswitch rather than forming a loop that interacts with the bound nucleotide. Finally, switch 2 is labile in the repressed state and does not interact strongly with the core. The N-terminus is myristoylated, such that if the N-terminal helix transiently dissociates from the core of the protein it can interact with appropriate membranes. Membrane interaction maintains the dissociation of the helix from the core, enhances nucleotide switching and ensures the correct localization of Arf function.

**Figure 1.**
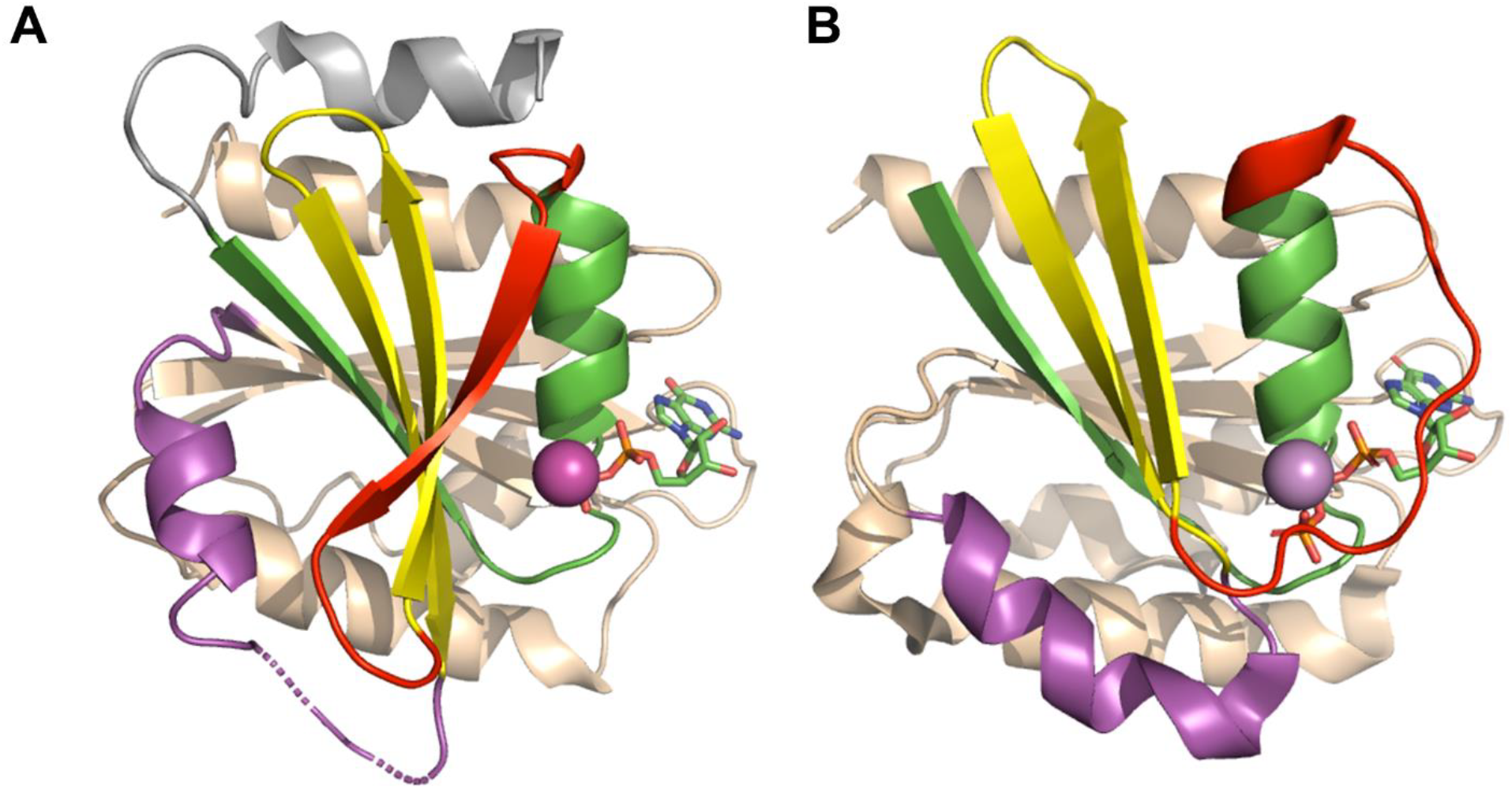
Structure of the GDP-bound and GTP bound states of Arf1. The structure of the GDP-bound, inactive state was taken from (14) (pdb:1hur) while that of the GTP-bound active state was taken from (30) (pdb:1j2j) in which the peptide from the GGA1 GAP is not shown. The first 17 residues including the N-terminal helix in the GDP-bound structure (A) are colored in grey, as they are not present in the Arf1Δ17 construct studied here. The rest of the N-terminal region is colored in green, switch 1 in red, the interswitch in yellow and switch 2 in magenta. The ligands (GDP in A and GTP in B) are CPK colored sticks. The C-terminal half of the protein is colored in wheat. The magnesium ion is light magenta. Coloring is the same for the GTP-bound form (B) except that the N-terminal 17 residues are not apparent in the structure. In the GTP-bound state (B) switch 1 (red) interacts with the bound GTP rather than forming a final strand of the central β-sheet as in (A), the interswitch β-hairpin is upshifted in register by two residues and switch 2 is more ordered and interacts more extensively with the core of the protein.

Structurally, the N-terminal helix in Arfs occupies the space that the interswitch must move into during the switch transition. This sterically represses the switch and also confines activity to the membrane. Indeed, spontaneous exchange is too slow to measure for full-length Arf1, but becomes readily observable, albeit not extremely rapid, if the first 17 residues encompassing the N-terminal helix are deleted, Arf1Δ17 (16), confirming repression of switching by this region of the protein. This front-to-back allosteric communication has been discussed previously (17). Note that here we will refer to it as top-down, given the orientation in which we present the protein structure (Figure 1A), and because we have previously reported a distinct allosteric pathway implicating communication between elements of the C-terminal half, on the backside of the protein, with the switch region, and which we termed back-to-front (18). Thus, while the structural role of the N-terminal helix in Arf protein function is well understood, the energetic consequences of its dissociation during the switch remain to be determined. Here we use the Arf1Δ17 construct to establish the energetic role of this unique feature of the Arf family of small GTPases.

## Materials and Methods

### Protein Production and Purification

Arf1Δ17 was overexpressed in E. coli BL21(DE3) in LB or ^15^N enriched minimal media as for FLArf1(19) a C-terminal His-Tag fusion using the pET-24b plasmid bearing a carboxypeptidase-blocking arginine residue immediately C-terminal to the Arf1Δ17 sequence. The C-terminal His-Tag was cleaved with 5 µL of ∼0.66 mM Carboxypeptidase A from bovine pancreas (Sigma-Aldrich) for every 1 mL of ∼0.40 mM of Arf1Δ17 in 50 mM Tris, 1 mM MgCl_2_, and 150 mM NaCl overnight at pH 7.5. Cleaved Arf1Δ17 was further purified and buffer exchanged to 50 mM bis-Tris, 150 mM NaCl, 1 mM MgCl2, and 5 mM DTT at pH 6.5 using a HiLoad™ 16/600 S75 Superdex pg (Cytiva). Purified Arf1Δ17 was loaded with GDP as previously described (20).

### High Pressure NMR

Amide backbone assignments for Arf1Δ17 were obtained from Buosi et al. (20). HP NMR was carried out as previously described (21) using the ceramic tube developed by Peterson and Wand (22) (Daedalus Innovations, Aston, PA). Detailed protocols for HP NMR experiments (^1^H-^15^N HSQC, and ^31^P NMR) can be found in the SI. Briefly, the chemical shifts associated with native protein HSQC backbone amide exhibit a distinct nonlinear response to pressure perturbation when lower volume excited states within the native-state manifold are populated, while linear responses to pressure result from simple compression. The linear and non-linear coefficients, *b*_*i*_ (ppm/bar) and *c*_*i*_ (ppm/bar 2), of pressure, *p*, dependent ^1^H and ^15^N chemical shifts for native state Arf1Δ17-GDP were determined by fitting the pressure-dependent changes in chemical shifts to a second order polynomial.

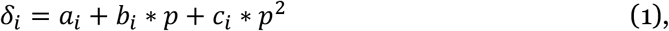

where *δ*_*i*_ is the chemical shift in ppm of *i*th residue.

The pressure-induced profiles for the loss of native state backbone amide ^1^H-^15^N HSQC peak intensities (I), were individually fitted to the following two-state model:

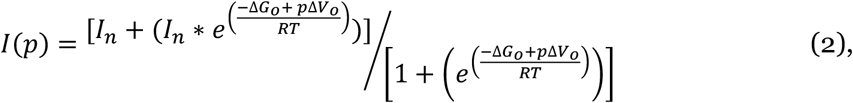

where *I*_*n*_ denotes the maximum peak intensity observed for the native state at atmospheric pressure, *p* is the pressure and *ΔG*_*o*_ and *ΔV*_*o*_ are the residue-specific apparent changes in the free energies at atmospheric pressure and molar volume. The 2-state model holds in this case because the amide is either in its native environment or it is not.

### High Pressure SAXS experiments

HP SAXS experiments were conducted on GDP loaded Arf1Δ17 at the ID7A beamline at the Cornell High-Energy Synchrotron source, CHESS at pH 6.5 as previously described (21). The 2D SAXS profiles were processed using BioXTAS RAW 2.1.4 package (23). The radius of gyration was determined using Guinier analysis and the pair distance distribution function (P(r)), computed with the GNOM package from the ATSAS 3.0.04-2 suite, integrated into the RAW software. (24). *High Pressure fluorescence measurements*

Time dependent HP fluorescence was carried out in a home-built HP optical cell using a modified ISS Koala fluorometer (ISS, Inc., Champaign, IL) as previously described (25), and an automated pump (Pressure Biosciences, Inc., Canton, MA). The exchange of GDP by GTP was measured using 20 µM of GDP loaded Arf1Δ17 at pH 6.5 with 50 M excess GTP at 33°C as previously described(26). A 120 s dead-time was required prior to data acquisition for sample loading and pressurization.

### Contact Maps and Cα-structure based molecular dynamics simulations

The native contacts in the protein were calculated from the crystal structure (1hur) with the N-terminal 17 residues removed using SMOG server (27). Using the contact information and the intensity from the HSQC data collected at 1, 1000, 1400 and 1800 bar, the fractional contact or probability of occurrence of all native contacts at each pressure (*FC*_*i*_*(p)*) was calculated as previously (28) as the geometric mean of the fractional intensities, *I*_*i*_*(p)/I*_*i*_*(atm)* and *I*_*j*_*(p)/I*_*j*_*(atm)*, at pressure, *p*, of the two residues, *i* and *j*, in the native contact:

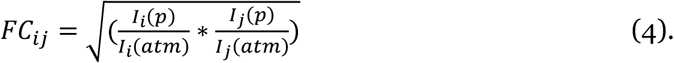

Fractional native contact plots are color-coded with red corresponding to the 100% native contact and blue to 0%.

## Results

### Arf1Δ17-GDP populates a heterogeneous molten globule ensemble under pressure

The ^1^H-^15^N amide HSQC spectrum of Arf1Δ17 was measured as a function of increasing pressure. The amide peaks shifted (Figure 2A) and their intensity decreased (Figure 2B) with increasing pressure, but no new peaks appeared. We have shown previously for full length Arf1 (FLArf1) that this is due to the pressure-induced population of a molten globule ensemble that undergoes conformational exchange on the NMR timescale (21). For Arf1Δ17, as for FLArf1, the midpoints of the pressure-dependent amide peak intensity decreases were observed over a broad range of pressures, <250 bar - >2000 bar (Figure 3). Moreover, amide peak intensity for residues in the N-terminal half of the protein decreases at lower pressure than for residues in the C-terminal half. These observations indicate the population of a heterogeneous ensemble of excited states.

**Figure 2.**
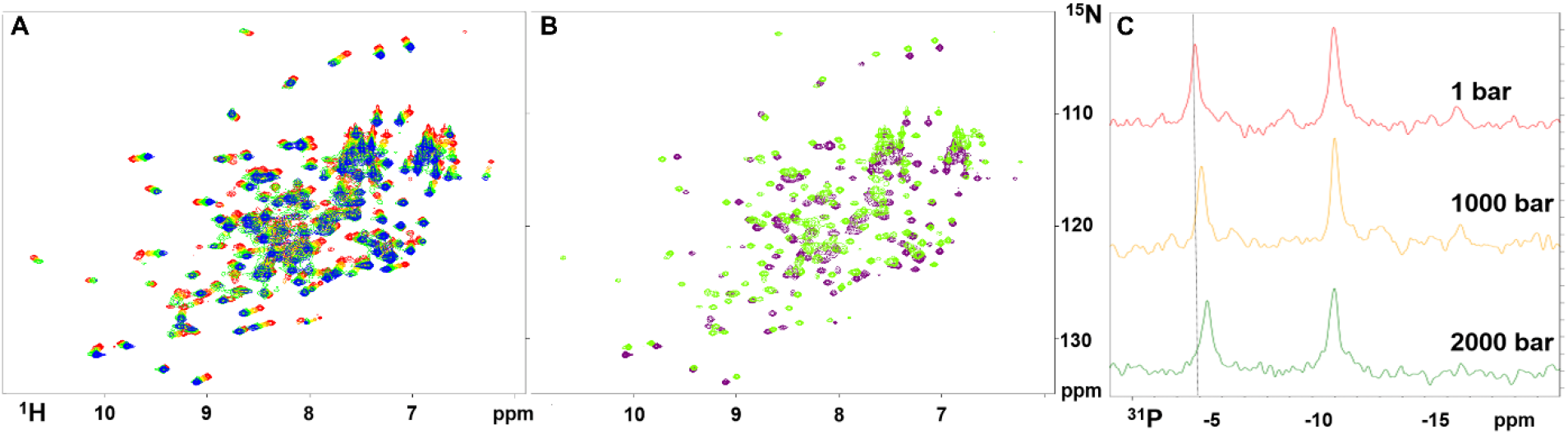
Pressure effects on Arf1Δ17-GDP NMR spectra. A) Pressure dependence of the ^1^H-^15^N HSQC spectrum of Arf1Δ17-GDP. Peaks shift from atmospheric pressure (red) to 2000 bar (purple) and decrease in intensity. B) Spectra for 1 bar (green) and 2000 bar (purple) highlighting the loss of peaks at high pressure. C) 1D ^31^P spectrum of Arf1Δ17-GDP as a function of pressure. The peak at ∼-4 ppm corresponds to the β-phosphate ^31^P resonance, while that at ∼-10 ppm corresponds to the bound α-phosphate ^31^P resonance of the bound GDP.

**Figure 3.**
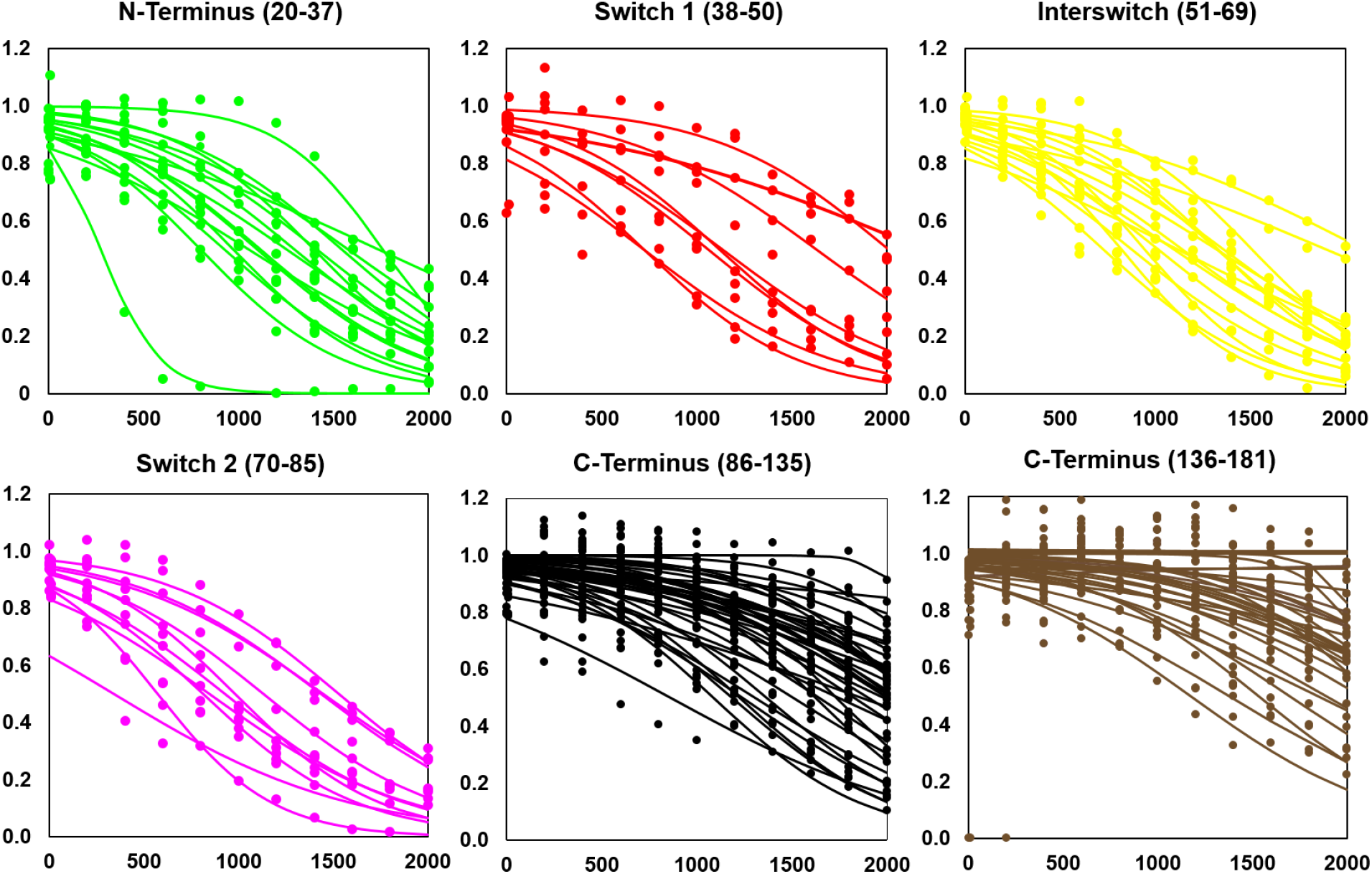
Pressure dependence of the amide peak intensities for most residues of Arf1Δ17-GDP. The different regions of the protein have been plotted separately for clarity. Color code corresponds to that in Figure 1 (N-terminal region, green; switch 1, red; interswitch, yellow; switch 2 magenta) except for the C-terminal half of the protein which has been separated in two parts, as indicated, for clarity and colored black and brown.

HP ^31^P NMR measurements of Arf1Δ17-GDP revealed that the nucleotide remained mostly bound, even at 2 kbar, although the chemical environment of the β-phosphate group was significantly impacted by pressure (Figure 2C), as previously observed for FLArf1 (21). In addition, at the highest pressure, 2000 bar, a small peak for the β-phosphate at - 6.85 ppm and a shoulder in the α-phosphate peak at -10.68 ppm were observed. The positions of the new peaks correspond to previously published values for free GDP•Mg^2+^ at high pressure (29), indicating a small degree dissociation may occur at 2000 bar.

We next turned to high pressure small angle x-ray scattering (HP-SAXS) to evaluate the effect of pressure on the global solution structure of Arf1Δ17. At atmospheric pressure, the pair distance distribution function revealed that Arf1Δ17 was folded and spherical, while high pressure led to subtle modifications in the P(r) plot (Figure 4A). Compared to FLArf1, the P(r) peak is shifted very slightly to smaller values and the extension is less apparent (Figure 4B). In the Kratky plots are normalized for size, it can be seen that Arf1Δ17 (green) was less spherical than FLArf1 (red) at atmospheric pressure (Figure 4C). While FLArf1 is compressed by pressure at this pH (6.5), Arf1Δ17 exhibited further deviation from a spherical shape at high pressure (purple), and the radius of gyration, R_g_, increased by ∼1Å (Figure 4D).

**Figure 4.**
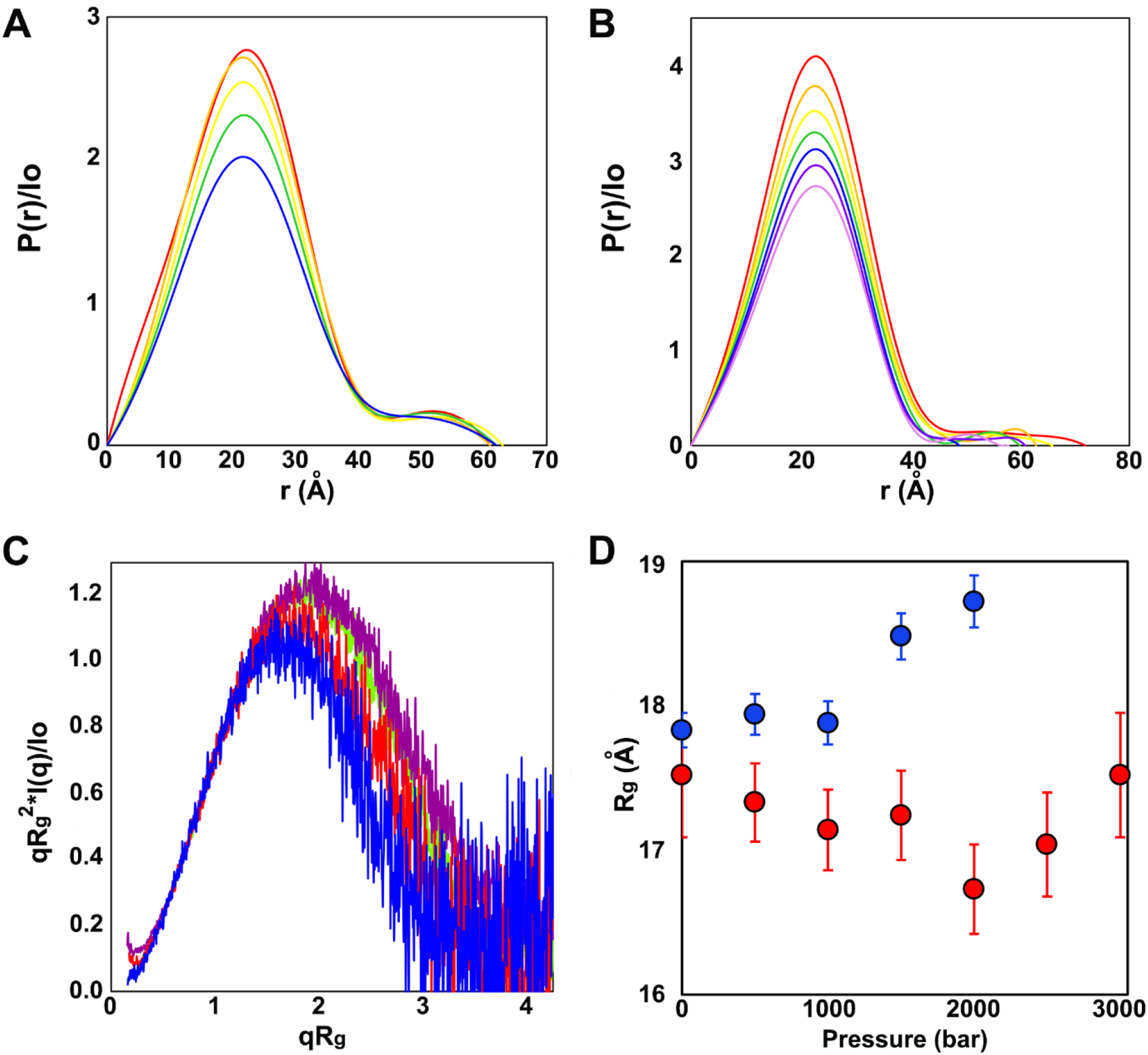
HP-SAXS of Arf1Δ17. A) P(r) pair distance distributions, respectively, of Arf1Δ17 at atmospheric pressure (red,) 500 bar (orange), 1000 bar (yellow), 1500 bar (green) and 2000 bar (blue); B) P(r) pair distance distributions, respectively, of FLArf1 at atmospheric pressure (red,) 500 bar (orange), 1000 bar (yellow), 1500 bar (green), 2000 bar (blue), 2500 bar (purple) and 3000 bar (pink); C) normalized Kratky plots of Arf1Δ17 at atmospheric pressure (green) and 2000 bar (purple) and of FLArf1 at atmospheric pressure (red) and 3000 bar (blue); D) Rg values vs pressure calculated from Guinier analysis of Arf1Δ17 (blue) and FLArf1 (red).

### Arf1Δ17-GDP is globally less stable than FLArf1-GDP

Fits of the pressure-dependent amide peak intensity loss profiles in Figure 3 to a two-state model yielded apparent residue-specific local free energy differences, *ΔG*_*app*_, between the native and excited states (Table S1). The N-to C-terminal stability gradient observed for FLArf1 (21) was conserved in Arf1Δ17-GDP (Figure 5A,B), with residues in the C-terminal half of the protein significantly more stable than those in the N-terminal, switch containing half. In contrast, the local stability of the majority of residues in Arf1Δ17-GDP was lower than for FLArf1 (21) (Figure 5C,D, Table S2). This global decrease in stability upon deletion of the N-terminal helix was distributed evenly across the protein (Figure 5C). Only 11 residues in Arf1Δ17-GDP exhibited a stability that was more than 0.5 kcal/mole more stable than FLArf1 (Table S1) (L25, V56, V68, G69, R97, V100, D141, S162, D164, Y154 and L173), many of which were among the least stable residues in FLArf1. No residue was found to be more than 1 kcal/mol more stable in the deletion mutant. While removing the N-terminal helix destabilizes the protein across its structure, there are some minor compensating effects.

**Figure 5.**
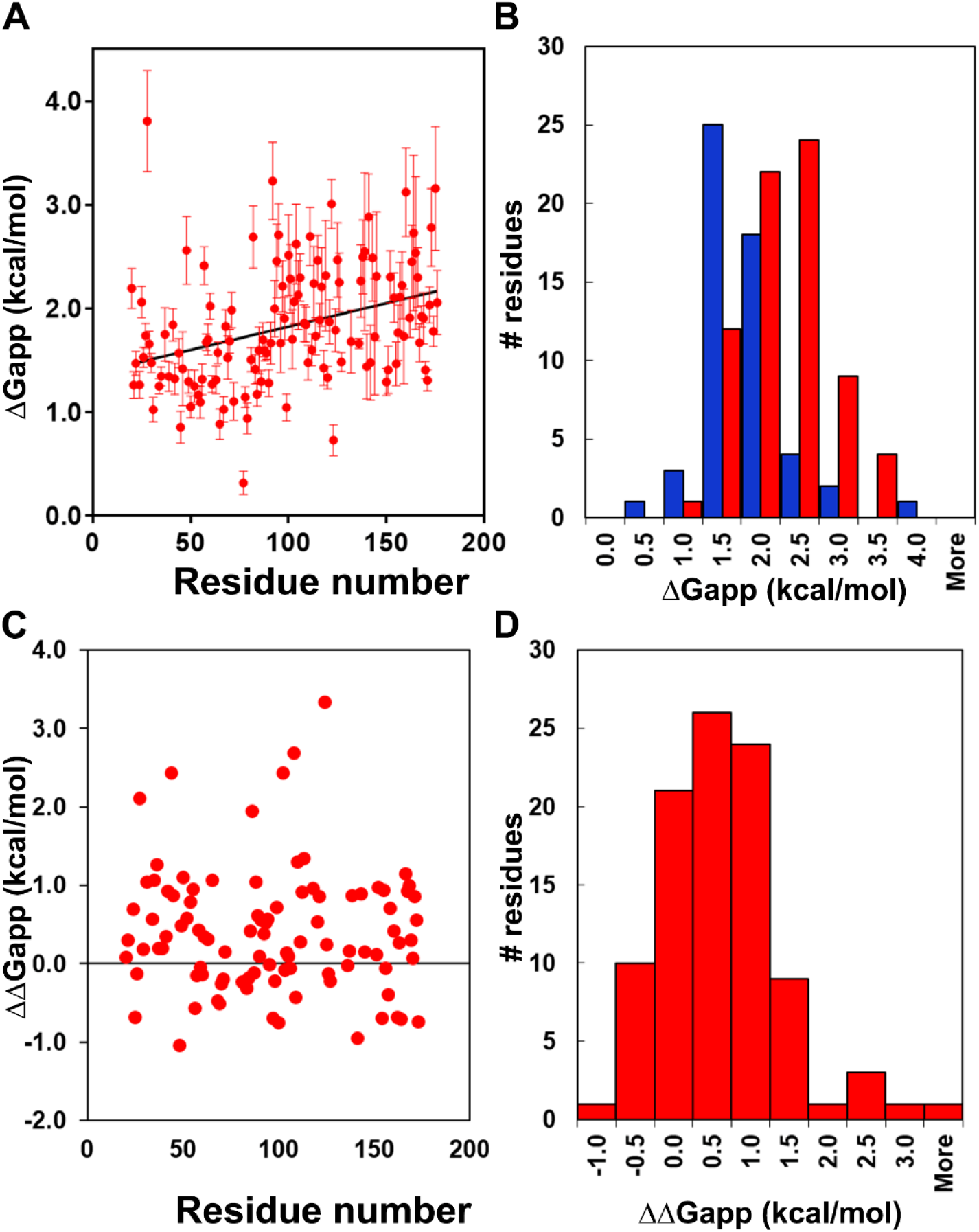
Local stability analysis for Arf1Δ17 from pressure dependence of NMR amide peak intensity. A) Residue-specific local *ΔG*_*app*_ values obtained from fits of the profiles in Figure 3. Histogram of the residue-specific local *ΔG*_*app*_ values for the N-terminal (blue) and C-terminal (red) halves of Arf1Δ17. C) Residue-specific *ΔΔG*_*ap*p_ values between FLArf1 and Arf1Δ17. D) Histogram of the residue-specific local *ΔΔG*_*app*_ values between FLArf1 and Arf1Δ17. Residue-specific local *ΔG*_*app*_ values for FLArf1 were taken from (21).

To provide a visual representation of the loss of native state under pressure, we used the pressure-dependent amid intensity profiles to calculate fractional native contacts at each pressure as the geometric mean of the fractional intensity of the two residues in a given native contact (Figure 6). As for FLArf1, the FCM for Arf1Δ17 at 1400 and 1800 bar highlight the bipartite nature of the stability distribution across the sequence. Also apparent from the FCM is the overall lower stability of Arf1Δ17 compared to FLArf1.

**Figure 6.**
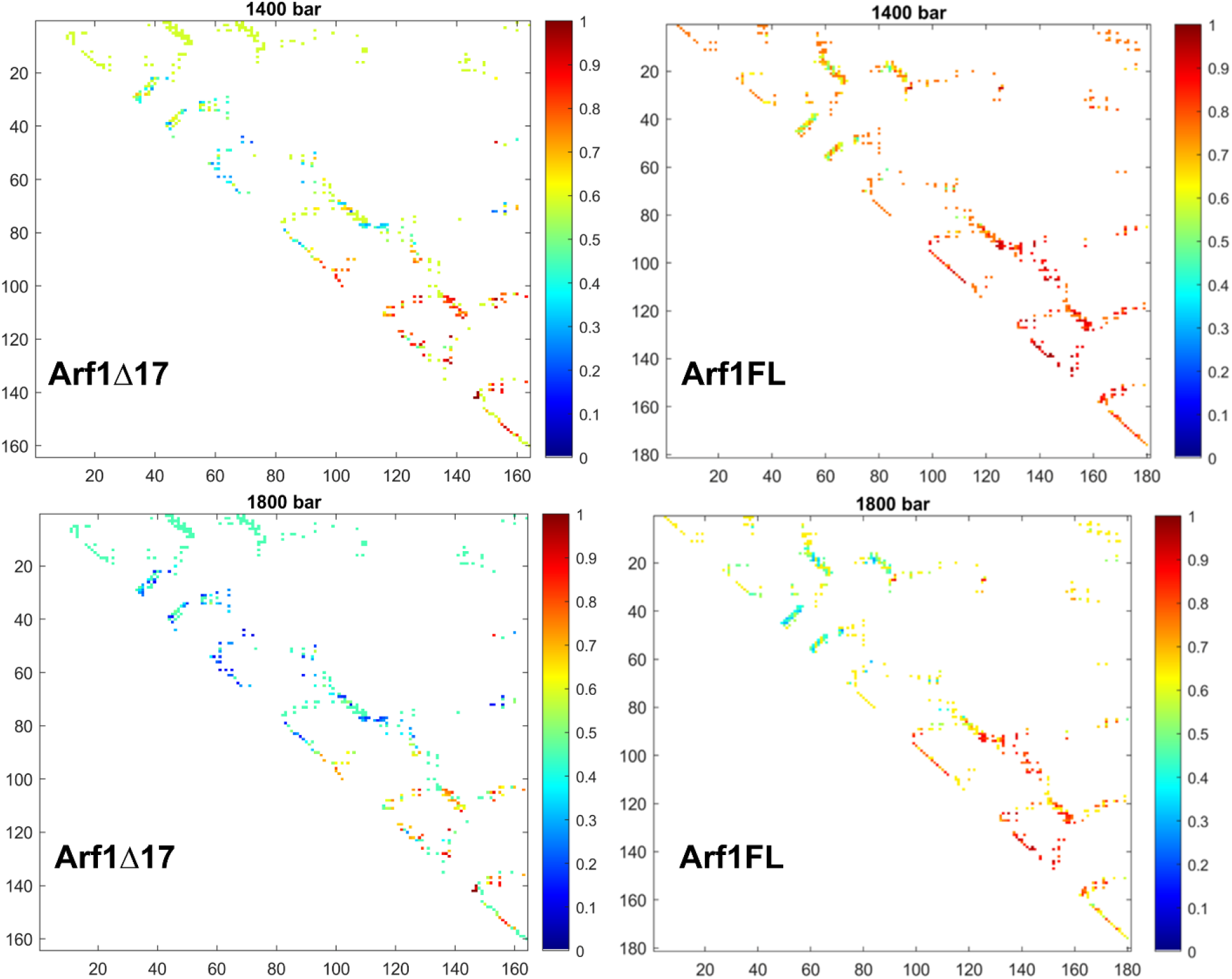
Fractional contact maps of Arf1Δ17 compared to FLArf1. Maps are shown for NMR data at 1400 and 1800 bar, as indicated. Fractional contacts are calculated as the geometric mean of the fractional intensities of the two residues in a given native contact at a given pressure, as indicated in the Materials and Methods section. Fractional contact maps for FLArf1 were replotted from (21).

## Discussion

Nucleotide switching in the Arf (and Arf-like) subfamily of small GTPases differs from all other small GTPases of the Ras superfamily (17). Most Arf family members exhibit a shorter interswitch hairpin and harbor an N-terminal helical extension that serves to maintain the protein in a retracted, inactive conformation when bound by GDP. To be activated, the proteins undergo an interswitch toggle, which requires dissociation of the N-terminal helix to allow for the interswitch to shift up in register by 2 residues. Switch 1 dissociates from the central β-sheet and moves several Å to interact with the triphosphate form of the nucleotide, while switch 2 also moves closer to the nucleotide. The structural mechanisms of this bipartite allosteric control, requiring on the one hand, GEFs to promote the interswitch toggle and, on the other, the membrane for N-terminal helix displacement, have been understood for some time (17). Here we have established the energetic mechanisms for repression by the N-terminal extension of Arf family proteins. Rather than a directional repression from the N-terminus to the switch, we find that the N-terminal helix stabilizes the protein across the whole structure. The previously reported (21) underlying N-to C-terminal stability gradient is maintained in the Arf1Δ17 N-terminal deletion variant, but the majority of residues are locally destabilized.

We recently demonstrated that the Arf family GTPases present an additional allosteric mechanism that controls their spontaneous switching rates(18). This mechanism implicates communication from elements in the C-terminal half to the switch region. Faster switching is imparted by lower local stability in the C-terminal regions, in particular in helix α5, where several substitutions relative to Arf1 are found. Moreover, strong, long distance (non-contact) evolutionary covariance between residues in the C-terminal half of a very large (>44,000) multiple sequence alignment of these homologous proteins suggests that this mechanism of switching control via local stability variations in the C-terminal half of the protein, distant from the switch, may be conserved across the entire Ras family of small GTPases. The present results suggest that the N-terminal helix acts to reinforce these C-terminal allosteric constraints, in addition to those that implicate the switch region directly and that are specific to the Arf family. The present results underscore the insight to be gained by mapping local protein energetics by coupling pressure perturbation with NMR spectroscopy. This approach could lead to new understanding of sequence-function relationships in a variety of important protein systems. In the future, coupling pressure perturbation with H/D XMS could extend the possibilities for gaining similar insights to much larger proteins and protein complexes.

## Author Contributions

Acquired data: EVP, TK, NH,

Analyzed data: EVP, TK, NH, SAM, CAR

Interpreted results: EVP, TK, SAM, JC, CAR

Wrote the paper: EVP, JC, CAR

## Acknowledgements

The authors would like to acknowledge Estella Yee of the BioSAXS beamline at MacCHESS for assistance in sample loading. The work was funded by a grant from the National Institutes of Health, GM 137766 to CAR and grants from the Fondation pour la Recherche Médicale (grant EQU202003010344) and the French Academy of Sciences (Grand Prix Emile Jungfleich, 2019) to J.C. The Center for High Energy X-ray Science (CHEXS) is supported by the NSF award DMR-2342336, and the MacCHESS resource is supported by NIGMS award 1-P30-GM124166-01A1. The Rensselaer Polytechnic Institute NMR core facility acknowledges National Institute of Standards and Technology grant 60NANB22D167 and National Institute of Health Grant 1S10OD 030482-01.

## Figures

